# Simulations suggest a scaffolding mechanism of membrane deformation by the caveolin 8S complex

**DOI:** 10.1101/2023.04.20.537670

**Authors:** Sayyid Yobhel Vasquez Rodriguez, Themis Lazaridis

## Abstract

Caveolins form complexes of various sizes that deform membranes into polyhedral shapes. However, the recent structure of the 8S complex was disk-like with a flat membrane-binding surface. How can a flat complex deform membranes into nonplanar structures? Molecular dynamics simulations revealed that the 8S complex rapidly takes the form of a suction cup. Simulations on implicit membrane vesicles determined that binding is stronger when E140 gets protonated. In that case, the complex binds much more strongly to 5 and 10-nm radius vesicles. A concave membrane binding surface readily explains the membrane-deforming ability of caveolins by direct scaffolding. We propose that the 8S complex sits at the vertices of the caveolar polyhedra, rather than at the center of the polyhedral faces.

**Significance:** Caveolae are membrane invaginations that play important roles in eukaryotic cells. Crucial for their formation is the caveolin protein family, but the mechanism by which these proteins deform membranes is unclear. Molecular dynamics simulations revealed that a recently determined structure changes shape from flat to concave. With one titratable residue protonated, this structure binds more strongly to spherical membranes, thus providing a possible mechanism for the membrane-deforming ability of caveolins.

## INTRODUCTION

Caveolae are invaginations in the plasma membrane of mammalian cells that have a distinct lipid composition enriched in cholesterol and glycosphingolipids (1). Their functions may include resistance to mechanical stress, mechanosensation, regulation of signaling, cholesterol homeostasis, trafficking, and endocytosis (2). Two proteins are essential for their formation: caveolins (3) and cavins (4). In humans there are three isoforms of caveolin and four isoforms of cavin. Caveolins by themselves have the capacity to deform the plasma membrane (5, 6) but formation of canonical caveolae also requires cavins.

Caveolin-1, the most prevalent isoform, has 178 residues organized in several domains: an unstructured N-terminal domain, the Oligomerization Domain (61-101) which includes the Scaffolding Domain (SD, 82-101), the so-called Intramembrane Domain (IMD, 101-134), and the C-terminal (Ct) domain. Caveolin was classified as an integral membrane protein based on its strong attachment to membranes (7). This and its topology (both termini in the cytoplasm) led to the proposal that the IMD inserts as a helical hairpin and acts as a wedge to deform the membrane (8). The monomeric IMD domain in detergent adopted a helix-break-helix structure (9). On the basis of this hypothesis, several theoretical studies explored the properties of the membrane embedded hairpin (10–13).

However, a recent cryoEM structure of the 8S complex showed a picture that was more reminiscent of a peripheral membrane protein (14). The complex contained 11 protomers with the SD, the IMD, and much of the Ct domain helical, ending in an unusual beta barrel with hydrophobic interior formed by the C-terminal ends. The paradox with this fascinating structure is that it presents a flat membrane binding surface, which was proposed to be deeply buried into the membrane so as to occupy one leaflet of the bilayer (14). It was also proposed to be located at and stabilize the flat faces of the observed polyhedra (15). But then, what causes the membrane bending in the absence of cavins?

In an attempt to confirm the proposed location in the membrane, we conducted molecular dynamics (MD) simulations of the 8S in an implicit solvent membrane and found that the complex rapidly took a conical shape with a concave membrane binding surface. An all-atom simulation in solution also showed the same shape change. Attachment to membranes was especially strong with protonation of the unique charged residue E140 on the hydrophobic face. We then used an implicit solvent model for curved membranes to calculate the binding energy to vesicles of different radius and found the strongest binding to vesicles of 5 and 10 nm radius. This finding leads to an alternative model of membrane incorporation and offers a clear mechanism for membrane curvature generation by caveolins.

## METHODS

The starting point for this work was the recent high resolution cryo-EM structure of the cav1 8S complex (pdb code 7SC0) (14). Residues 1-48 and 178 were not visible in this structure and were omitted. Implicit solvent simulations in solution were conducted using the effective energy function EEF1, which is a combination of the CHARMM 19 united atom force field (16) with a Gaussian solvent-exclusion solvation free energy term (17). The starting structure was minimized for 300 steps using the Adopted Basis Newton Raphson method and then subjected to microcanonical MD. The average temperature was kept at 300 K using periodic rescaling of velocities if necessary.

Simulations in solution were also carried out with an all-atom representation with the TIP3P water model and the CHARMM36 force field for the protein (18). Here, the complex was inserted into a rectangular box with sides 186.6, 186.6, and 93.3Å filled with TIP3P water. Water molecules within 2.6 Å from protein heavy atoms were deleted. The final system contained 99,541 water molecules and 33 K^+^ ions that were added to neutralize the unit cell, bringing the total number of atoms to about 317,500. With harmonic constraints on the protein (force constant 5 kcal/mol.Å), the system was minimized for 5000 Steepest Descent steps and 300 Adopted Basis Newton Raphson steps. Then the constraints were released and the MD simulation started. The nonbonded cutoff was 12 Å and Particle Mesh Ewald was used for long range electrostatics. The simulation was run at constant temperature of 303 K using the Hoover thermostat and constant pressure of 1 atm using the Langevin piston barostat. This simulation crashed at 7.4 ns when the complex rotated and touched the unit cell boundary. We analyze the data up to 7.35 ns.

The membrane simulations used the IMM1 model (19) and its extension to curved membranes (20), which we previously used to study ESCRT-III snf7 (21) and IBAR domains (22). IMM1 is an extension of EEF1 to heterogeneous membrane-water systems. It models the membrane as a hydrophobic slab and uses a switching function that transitions smoothly from a non-polar to an aqueous environment. The headgroup area is not distinguished from the aqueous region, so specific interactions with the lipid headgroups are ignored. Only the headgroup charge is taken into account using the Gouy-Chapman model (23). Extension to curved membranes was achieved by changing the definition of relative depth z’ from |z|/(T/2), where T is the thickness of the membrane, to |r-R|/(T/2), where R is the radius of the vesicle or tube and r the radial position of an atom. The model also allows one to include the change in lateral pressure profile as the membrane bends, but for simplicity this feature was not used in the present simulations.

All MD simulations were carried out with the CHARMM software package (24). The implicit membrane simulations were run at 300 K with a 2-fs time step for 2 ns, a time that was found sufficient for convergence of the structural change in solution. Due to the lack of solvent friction, the true time-scale of implicit solvent simulations is longer than their nominal duration. Two replicate simulations with different initial velocities were run for the protonated E140 systems and the simulation in implicit water was extended to 25 ns to test convergence.

Membrane binding energies were estimated as average transfer energies, i.e. the difference in energy of the peptides on the membrane surface and the same conformation in bulk water. Here, for each snapshot from the simulation on the membrane, the protein is translated to the bulk and the energy change is calculated and then averaged over the trajectory. This allows us to reduce the statistical error, but is approximate, because it neglects possible changes in intramolecular energy. Conformational and translational entropy is also neglected in these calculations, but these contributions are likely to be very similar in binding to membranes of different curvature.

The shape of the complex was monitored by first orienting the complex with its first two principal moments on the xy plane and then measuring the difference between minimum and maximum x, y, and z coordinates (Δx=X_max_-X_min_, etc) as a function of simulation time. The energetic reasons for the observed conformational change were explored in the implicit water simulation by comparing the effective energy of the minimized experimental structure with that of the minimized structure after 5 ns dynamics. The energy was decomposed into various contributions, intrachain, interchain etc. using the CHARMM INTE command. Then, simulations were run using the CHARMM CONS RMSD command, constraining different sections of the chain each time to remain close to their original conformation. In these trials each protomer was constrained independently of the others.

## RESULTS

### Simulations in solution

Caveolin is constitutively attached to membranes. However, after we observed a large conformational change in implicit membrane simulations, we thought it useful to explore the conformational properties of the 8S complex in the absence of lipids or detergent. So, we started with a 25-ns MD simulation of the 8S complex in aqueous solution using the EEF1 implicit solvation model (17). The overall structure was well-maintained but the shape rapidly changed from flat to conical (Fig. 1). Fig. 2 shows the backbone RMS deviation of the entire complex from the initial structure and the extent of the complex in the x, y, and z directions when aligned with the xy plane. There is a rapid increase in the height (Δz) of the complex from 40 to about 60 Å. The diameter does not change much, except for a transient increase in the beginning and some loss of symmetry. The changes converge in 2-3 ns. Two replicate 5-ns simulations with different initial velocities gave very similar results (one gave final backbone RMSD 9.6 Å and Δz = 53.9 Å and the other 11.1 Å and 62.1 Å, respectively).

**Fig. 1.**
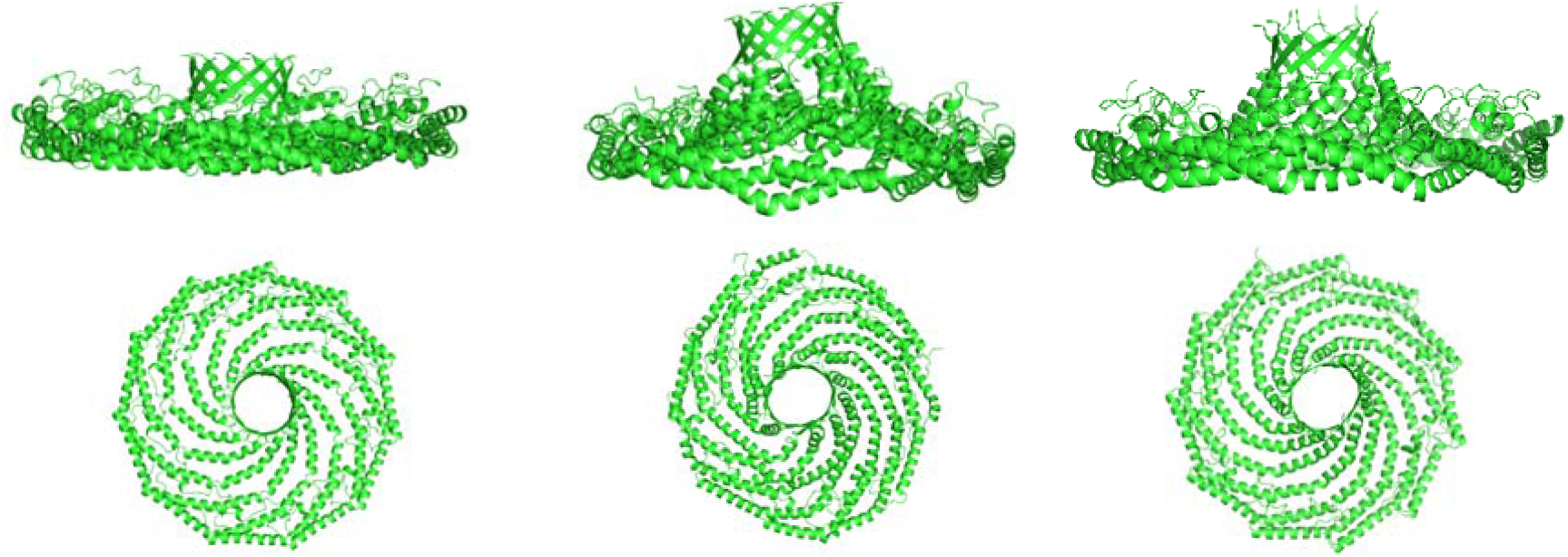
Structure of 8S before simulation (left), after 5-ns of MD simulation in implicit solvent (middle), and after a 7.35-ns MD simulation in explicit solvent (right). Side views above and top views below.

**Fig. 2.**
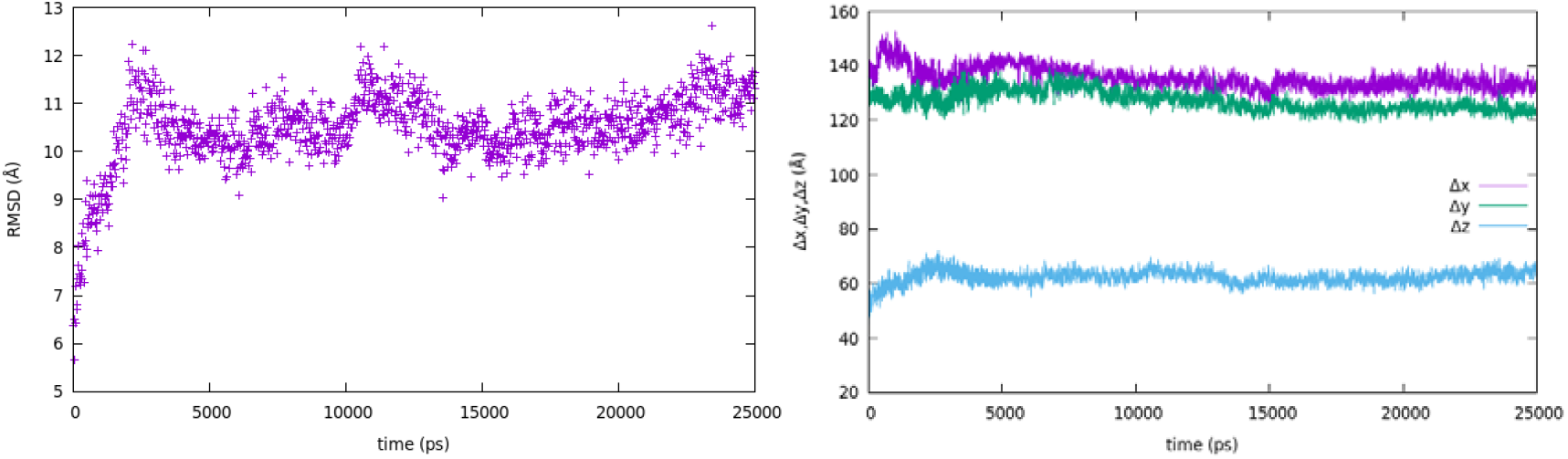
25-ns implicit solvent simulation in solution. a) Backbone RMS deviation of the entire complex with respect to the initial structure, b) Spatial extent of the complex (the values for the cryoEM structure are 137.4, 137.4, 40.1 Å)

To check the validity of this approximate implicit solvent energy function the complex was also simulated in all-atom aqueous environment. The size of the system restricted us to only 7.35 ns but that was sufficient to show a similar structural change as the one observed in the EEF1 simulations, except it happens more slowly (Fig. 3). The backbone RMSD reaches just above 7 Å (compared to 10 Å with EEF1) and keeps slowly increasing. The width of the complex increased rapidly from 40.1 to about 53 Å within 0.5 ns and then more slowly to about 58 Å, again with increasing trend. The diameter decreased slightly to about 130 and 136 Å, with some symmetry lost as the simulation progressed.

**Fig. 3.**
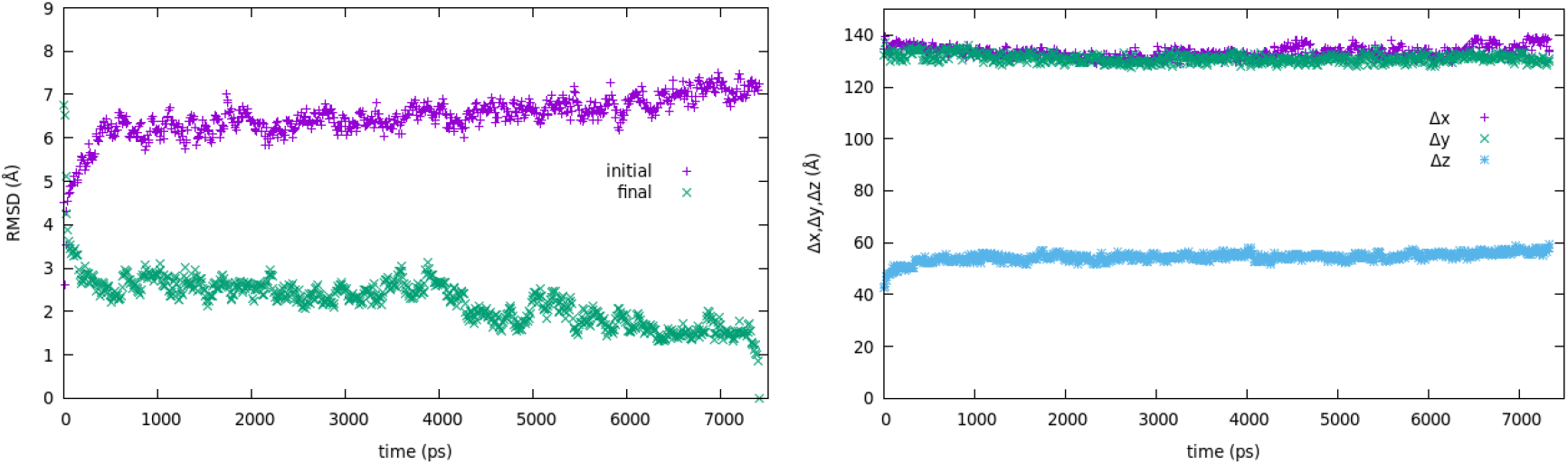
Backbone RMSD (a) and spatial extent (b) and as a function of time in a 7.35-ns all-atom simulation in water. In (a) the RMSD is calculated with respect to the initial or the final conformation.

### The cause of the observed conformational change

The reason for the conformational change is easier to investigate in the implicit solvent runs. We did that in two ways. First, the reduction in effective energy from the minimized experimental structure to the minimized structure after 5 ns was decomposed into contributions using chain (protomer) A as an example (Table 1). It was found that both intrachain and interchain interactions improve, but the intrachain energy change is larger and comes mainly from improvement within the backbone and in the side chain-backbone interactions. The largest contribution to the latter comes from the charged and polar side chains. The improvement in the backbone energy appears to arise from an increase in the number of H bonds. Using a cutoff of 3.5 Å and the HBOND facility in CHARMM, we calculated 168 H bonds for a protomer in the original structure vs. 187 in the MD-relaxed structure. The experimental structure exhibits several breaks or turns in the helical structure. In addition to three Pro at 110, 132, and 158, there is a π helix at 126-130 and a short 310 helix at 163-167. In the MD structure a break develops in helix α1 and the rest of the chain becomes more continuous (Fig. 4). The π-helix is maintained during MD.

**Table 1.** Contributions to change in effective energy (kcal/mol) from experimental structure to the MD structure at 5 ns. A and B refer to two protomers, b=backbone, s=side chains, AsKRDE are the charged side chains of protomer A, AsFYW the aromatic side chains of A, and so on.

| | cryoEM | MD | $\Delta W$ |
| --- | --- | --- | --- |
| Total W per protomer | -1400 | -1606 | -206 |
| intra-A | -1186 | -1315 | -129 |
| A – B | -199 | -285 | -86 |
| intra-Ab | -1102 | -1145 | -43 |
| intra-As | -37 | -40 | -3 |
| Ab – As | -70 | -155 | -85 |
| Ab – AsKRDE | -50 | -86 | -36 |
| Ab – AsSTNQ | -31 | -67 | -36 |
| Ab – AsALIV | +31 | +27 | -4 |
| Ab – AsCMP | +2 | -2 | -4 |
| Ab – AsFYW | -23 | -27 | -4 |
| AsKRDE – BsKRDE | -11 | -11 | 0 |

**Fig. 4.**
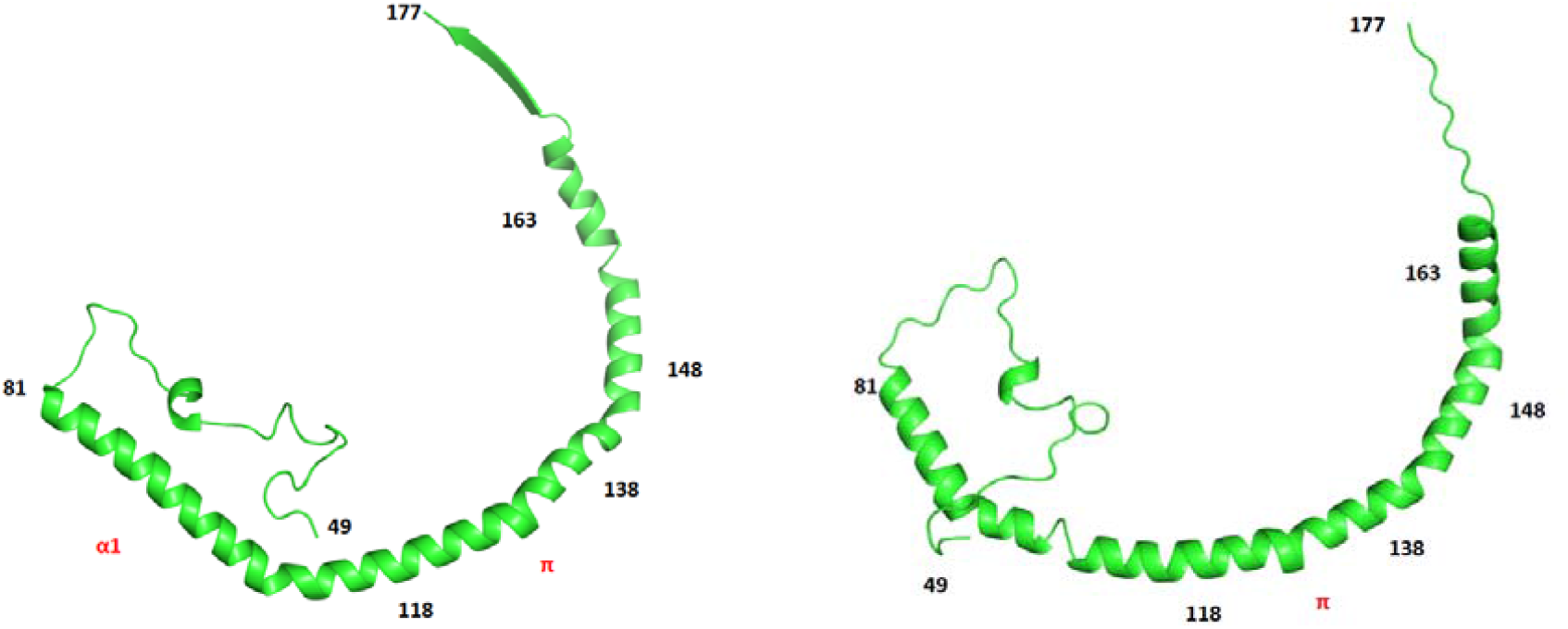
The protomer in the experimental structure and after 5-ns implicit solvent MD

To pinpoint which section of the backbone is responsible for the conformational change, we performed brief (50-ps) simulations with parts of the structure constrained. The segments were selected to span specific turns or breaks in the helix. Table 2 shows that constraining the backbone of residues 81-177 of each chain separately prevents the shape change. Constraining shorter segments of the backbone also prevents the change, as long as they are at least 20-25 residues long. So, it is the change in backbone structure of each chain to one that is more continuous and with more H bonds that causes the rise in the height of the complex.

**Table 2.** Backbone RMSD of the entire 8S complex and height (spatial extent in the z direction) upon a 50-ps implicit solvent MD constraining different segments of the chain. Each chain (protomer) is constrained independently of the others.

| Segment Constrained | RMSD | $\Delta z$ |
| --- | --- | --- |
| None | 7.4 | 54.3 |
| 49-177 (entire protomer) | 0.4 | 42.7 |
| 81-177 | 1.8 | 42.2 |
| 81-138 | 3.0 | 45.1 |
| 81-118 | 3.7 | 45.2 |
| 118-148 | 4.2 | 44.5 |
| 118-138 | 5.3 | 43.5 |
| 138-163 | 3.7 | 43.1 |
| 138-148 | 7.7 | 58.0 |
| 148-163 | 9.6 | 63.9 |

### Simulations on implicit flat and curved membranes

To examine the interaction of 8S with membranes we used the IMM1 implicit membrane model (19) and its extension to curved membranes (20). The complex was placed with its flat hydrophobic face at the interface or buried up to midplane of an implicit membrane. The conformational change observed in solution also occurred on the membrane. In fact, it is even more rapid and transiently overshoots in z extent to 69 Å before it relaxes to values similar as in solution (Fig. 5a). The complex binds strongly and the outer ridge buries quite deeply in the membrane (Fig. 5b), as predicted (14). Most deeply buried is F107, which reaches 4.5 Å from the center of the membrane. However, the central region of the disk does not insert. The presence of negative surface charge on the flat membrane did not improve the binding but instead weakened it somewhat when E140 was ionized (Table 3). The basic residues in 8S are on top of the complex and far from the membrane surface, so they feel weakly the surface charge.

**Table 3.** Average transfer energies (kcal/mol) of caveolin 8S from water to the membrane surface. The middle column is for ionized E140 and the right column for protonated E140. All membranes are neutral except for the last entry.

| Radius (nm) | $\langle\Delta W\rangle$ (E140<sup>-</sup>) | $\langle\Delta W\rangle$ (E140H) |
| --- | --- | --- |
| 5 | $-92 \pm 15$ | $-332 \pm 37$ |
| 10 | $-224 \pm 43$ | $-313 \pm 50$ |
| 25 | $-225 \pm 14$ | $-257 \pm 11$ |
| 50 | $-230 \pm 13$ | $-262 \pm 9$ |
| Flat | $-217 \pm 14$ | $-248 \pm 7$ |
| Flat 30% anionic | $-205 \pm 11$ | $-260 \pm 6$ |

**Fig. 5.**
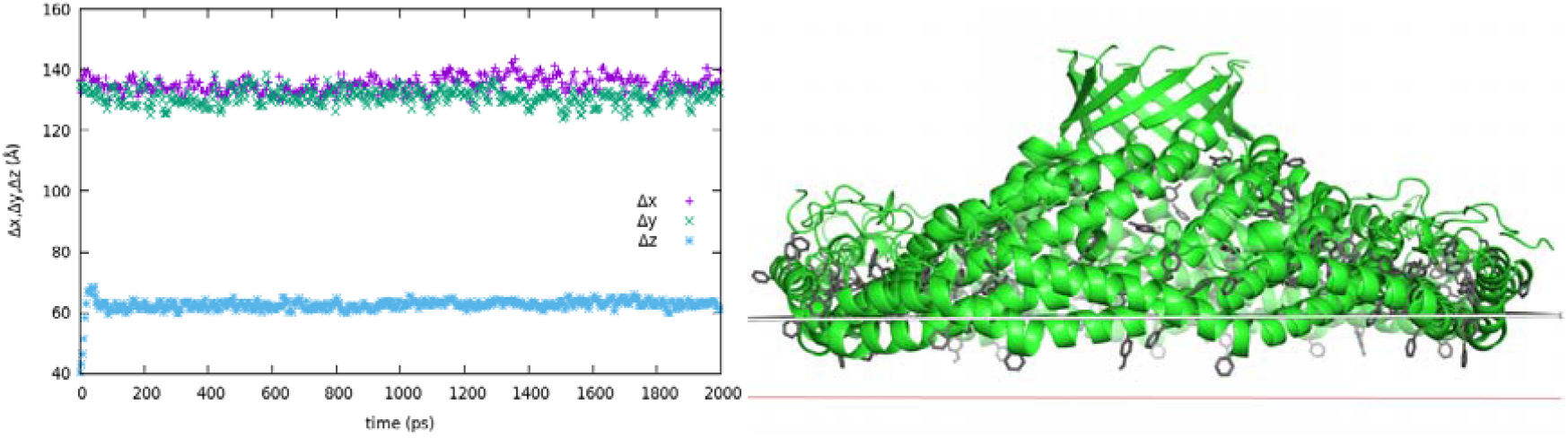
Spatial extent as a function of time (a) and conformation (b) of 8S on a flat, neutral implicit membrane. Side view showing membrane insertion. The grey line is the hydrophobic-hydrophilic boundary and the red line the bilayer midplane. Tyr and Phe side chains are shown in stick representation.

We noticed that the only charged residue on the membrane binding face is E140, which, especially in its ionic form, is repulsed by the membrane surface. Protonation of a Glu costs only 2.3RT(pK_a_-pH), i.e., assuming a standard pK_a_ and neutral pH, about 3.5 kcal/mol, or 38 kcal/mol for the entire complex. So, we also conducted simulations with E140 protonated, which improves the binding (Table 3).

The complex was then placed at vesicles of radius 50, 100, 250, and 500 Å starting from the final structure after MD in water and was simulated for a further 2 ns. Fig. 6 shows the final conformations on the different size vesicles and Table 3 shows the transfer energy from water to the membrane at vesicles of different radius and the flat membrane. For anionic E140 there is little change in binding energy going to curved membranes. The complex binds only at the ridge, and that does not change in the different vesicles. In fact, it binds worse at the smallest vesicle. When E140 is protonated, however, the complex binds using a larger portion of its hydrophobic surface. In this case, curvature makes significant difference and binding is very tight on the smaller vesicles. The difference in binding energy between protonated and unprotonated E140 at high curvatures is sufficient to “pay” the cost of protonation. Anionic charge on the flat membrane improves somewhat the binding when E140 is protonated but not when it is charged.

**Fig. 6.**
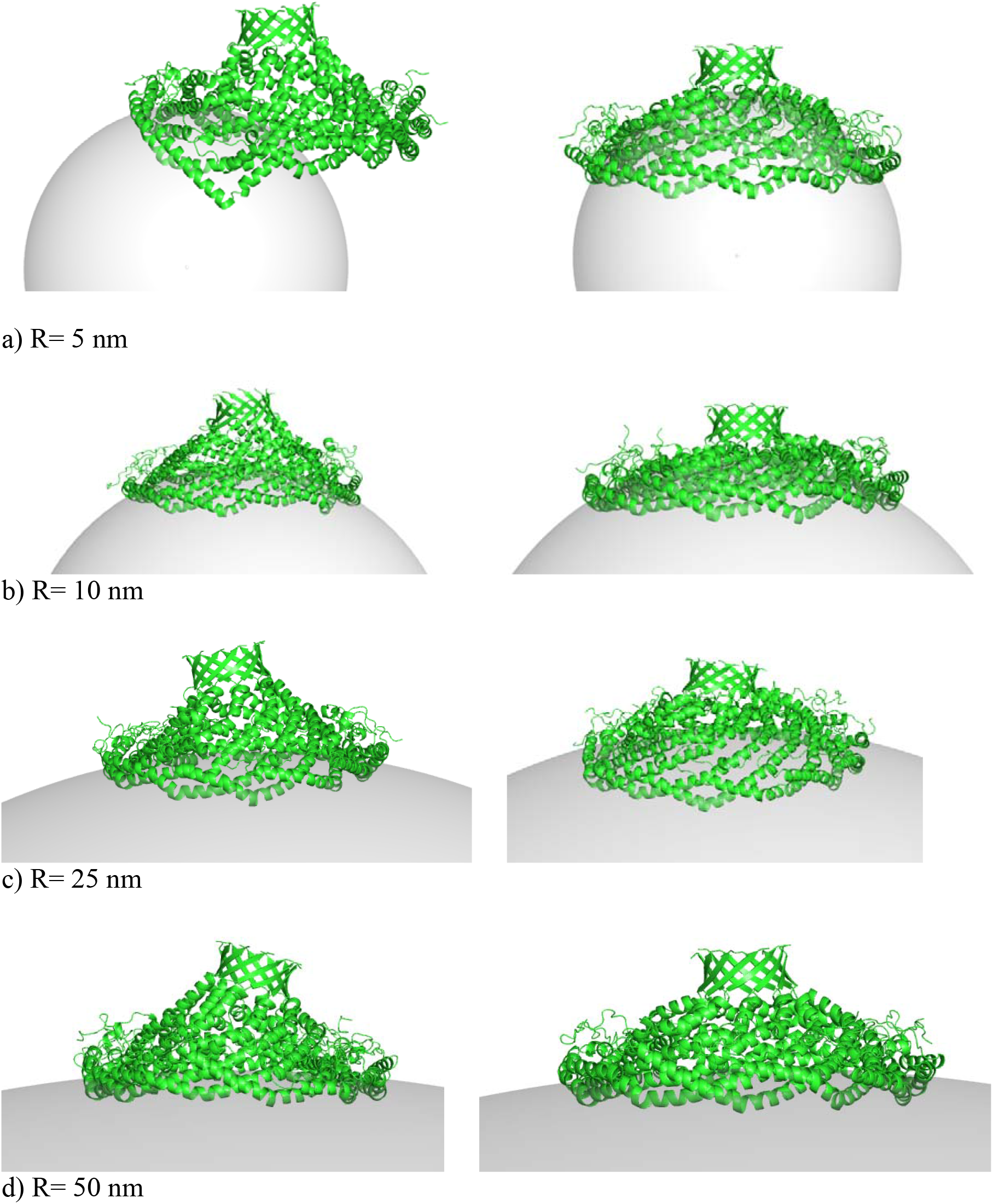
Conformation of 8S on vesicles of different radius. E140 is ionized on the left and protonated on the right. a) R=5 nm, b) R=10 nm, c) R=25 nm, d) R=50 nm.

To characterize the shape of the complex in the different vesicles we calculated the spatial extent of the complex (Table 4). The height of the complex (Δz) is smaller when E140 is protonated. With respect to vesicle size, Δz is smallest for the 10-nm vesicle. At that curvature the complex resembles most the cryoEM structure. Asymmetry is larger when E140 is ionized.

**Table 4.** Spatial extent of the complex on the different membranes in Å. The standard deviations are 1-4 Å. The values in the cryoEM structure are 137.4, 137.4, 40.1.

| Radius (nm) | $\Delta x, \Delta y, \Delta z$ (E140<sup>-</sup>) | $\Delta x, \Delta y, \Delta z$ (E140-H) |
| --- | --- | --- |
| 5 | 133.6, 127.2, 71.3 | 134.3, 130.2, 63.6 |
| 10 | 137.1, 128.0, 67.9 | 140.3, 135.1, 52.4 |
| 25 | 144.0, 128.9, 64.0 | 136.0, 130.7, 59.4 |
| 50 | 141.9, 130.1, 63.7 | 135.9, 130.7, 59.7 |
| Flat | 143.7, 128.5, 63.2 | 134.4, 131.1, 56.1 |
| Water | 138.8, 128.8, 61.9 | 134.5, 125.5, 60.4 |

To test the reproducibility of the results we performed two additional replicates for the E140H runs starting with different random velocities. The resulting <ΔW> values were within the error bar reported in table 3 and the spatial extents similar to those in Table 4. To also test the sensitivity to starting conformation, we repeated the simulations starting from either 5 Å closer to or 5 Å further from the membrane. Again, the results were within the uncertainty, except that for the +5 Å simulations at 5 and 10 nm it took some time for the complex to bind to the membrane and that time had to be excluded from the averaging to obtain congruent results.

### Simple pK_a_ calculations for E140

To confirm the role of E140, we made simple pK_a_ estimates using the implicit solvent model used for the simulations. For these calculations we take each configuration of interest, minimize the energy with E140^-^ for 300 steps, record the energy (W1), add a proton to generate E140H, and record the energy again (W2). So, ΔW=W2-W1 is the difference in effective energy upon protonating E140. This number in itself has no physical meaning, but the change in ΔW between the system of interest and a model system where a Glu is fully solvent exposed is related to the ΔpK_a_ according to ΔΔW = - 2.3 RT ΔpK_a_. As model system we use a 25-residue polyalanine helix with a Glu in the middle. Table 5 shows the ΔΔW and ΔpK_a_ values calculated for some of our structures.

**Table 5.** pK_a_ estimates for E140. ΔW is the change in energy upon protonating E. ΔΔW is the change in ΔW from that in the polyAla helix.

| | $\Delta W$ | $\Delta\Delta W$ | $\Delta pK_a$ | $pK_a$ |
| --- | --- | --- | --- | --- |
| E in polyAla helix | 15.9 | - | - | 4.3 * |
| E140 in cryoEM str in water | 13.8 | -2.1 | 1.6 | 5.9 |
| E140 in water, after MD | 15.0 | -0.9 | 0.7 | 5.0 |
| E140 in cryoEM, neutral mem | 10.7 | -5.2 | 3.8 | 8.1 |
| E140 in cryoEM, anionic mem | 8.9 | -7.0 | 5.1 | 9.4 |
| E140 at the E140H structure<br>after MD on R=5 nm vesicles | 8.7 | -7.2 | 5.2 | 9.5 |

\* The $pK_a$ of Glu in a polyAla helix is assumed to be the same as a Glu in a pentapeptide (25).

These simple estimates are consistent with our previous observations. We note that these have been obtained by single point energy calculations on minimized structures. More rigorous calculations, along with estimates of statistical error, can be obtained by MD sampling the configurational space under constraints. It also possible to calculate pH-dependent membrane binding energies using the present implicit solvation model (26).

## DISCUSSION

The structures resulting from the simulations reported here depart significantly from the recent cryoEM structure (14) as well as earlier lower resolution work (15, 27) that show the caveolin 8S complex to be a flat disk. This would normally lead us to discount the simulation results as artifacts of the imperfections of the energy functions. However, because they make more mechanistic sense than the experimental structures, it is worth entertaining the possibility that some experimental factor biases the experimental structures toward flatter structures. The biggest difference between the in silico and in vitro systems is the presence of large amounts of detergent in the latter (dodecyl maltoside (14, 27) or octyl glucoside (15)). Structural work seems to be quite sensitive to the type of detergent used (28). Detergent covers 60% of the surface area of the complex, including much of the central cavity (14). It is not immediately clear how the detergent would flatten the disk, given that detergents are usually mobile and pliable and can adapt to a given shape. We plan to investigate this in the future by including detergent in the simulations.

A recent article reported remarkable agreement between the caveolin 8S cryoEM structure and the predictions of Alphafold2 for caveolin monomers or oligomers (29). Interestingly, some predictions contains deviations similar to the ones observed in our simulations, namely, the upward movement of the beta barrel, a larger extent in the z direction, and some concave curvature on the membrane binding surface. Also interesting is that the predictions contain more continuous helices, something that also results from our MD simulations.

A residue with a profound effect on binding to curved membranes was found to be E140. This residue is also E in cav3 and V in cav2. It can be hydrophobic or polar in other species (30). To our knowledge, there has not been any experimental investigation of this residue and we can only speculate on its role. Is it there to moderate binding affinity and thus allow caveolae to dissolve in response to mechanical stress? Does it impart pH sensitivity? In any case, it is an obvious target for mutation. It should be noted that cav1 is palmitoylated at three nearby positions (133,143,156). These palmitoyl groups, which are not included in the present simulations or in the cryoEM structure, would certainly increase the attachment to membranes. Nevertheless, palmitoylation is not necessary for localization in caveolae (31).

Electron microscopy has shown that caveolae have a polyhedral shape and that cav1 forms the inner part of the coat and cavin 1 the outer layer, with filaments running parallel to the edges of the polyhedra (32). Heterologous caveolae, i.e. structures formed in *E. Coli* after heterologous expression of cav1, also have polyhedral shape (5, 33). A model for the structure of cavin oligomers has also been proposed (15). The flatness of 8S led to the proposal that it binds the flat faces of the polyhedra, buried quite deeply in the membrane by displacing one bilayer leaflet (14, 15). But then, what deforms the membrane in the absence of cavins? If 8S can sense and generate membrane curvature itself, it may actually occupy not the flat faces but the vertices of the polyhedra observed in EM. The connections between the vertices could perhaps be provided by the missing, disordered N-terminal domains which could adopt defined structure under the right conditions.

The common size of caveolae is 50-80 nm in diameter (34), and heterologous caveolae in *E. Coli* induced by cav1 alone are 45-50 nm (5). Here we find stronger binding to smaller vesicles of diameter 10 and 20 nm. Several comments can be made in regard to this discrepancy. First, the transfer energy that we compute is not the full free energy of the protein (it does not include any intramolecular distortion energy of the complex). Second, the diameter of a vesicle is determined by a balance between binding energy and membrane deformation energy, which tends to keep the membrane flat. This energy is also not included in our calculation. Finally, if the 8S truly occupy the vertices of the polyhedra, the local curvature there will likely be higher than the overall curvature of the caveolae.

The caveolin story seems to offer one more example in the membrane remodeling field where the monomer by itself is irrelevant. It is the properties of the oligomer that determine function. This has also been recently suggested for the septins, although their structure is not yet known (35). In previous work from this lab, the same was suggested for the IBAR domains (22), but is not yet experimentally confirmed. Scaffolding by an oligomer is thus a strong candidate mechanism for membrane curvature generation.

One significant limitation of the present computational approach is that the membranes are not deformable. Simulating the 8S complex in the presence of an all-atom bilayer is necessary to confirm the mechanism of action that we propose. Also interesting would be the simulation of the complex in the presence of detergent. These demanding simulations are in our immediate plans.

## Acknowledgments

We thank the National Science Foundation (MCB-1855942) for funding and Dr. Amedee Des Georges for useful discussions.

## Author contributions

TL conceived the project and wrote the article. SYVR conducted the simulations, analyzed the data, and prepared the figures with a small contribution from TL.

